# Overproduction of the AlgT sigma factor is lethal to mucoid *Pseudomonas aeruginosa*

**DOI:** 10.1101/2020.06.05.136770

**Authors:** Ashley R. Cross, Vishnu Raghuram, Zihuan Wang, Debayan Dey, Joanna B. Goldberg

**Affiliations:** Division of Pulmonary, Allergy & Immunology, Cystic Fibrosis and Sleep, Department of Pediatrics, Emory University School of Medicine, Atlanta, GA USA; Emory+Children’s Center for Cystic Fibrosis and Airway Disease Research, Emory University School of Medicine, Atlanta, GA USA; Department of Biochemistry, Emory University School of Medicine, Atlanta, GA USA

**Keywords:** *Pseudomonas*, gene regulation, sigma factor, proteolysis, lethality

## Abstract

*Pseudomonas aeruginosa* isolates from chronic lung infections often overproduce alginate, giving rise to the mucoid phenotype. Isolation of mucoid strains from chronic lung infections correlates with a poor patient outcome. The most common mutation that causes the mucoid phenotype is called *mucA22* and results in a truncated form of the anti-sigma factor MucA that is continuously subjected to proteolysis. When a functional MucA is absent, the cognate sigma factor, AlgT, is no longer sequestered and continuously transcribes the alginate biosynthesis operon leading to alginate overproduction. In this work, we report that in the absence of wild-type MucA, providing exogenous AlgT is toxic. This is intriguing since mucoid strains endogenously possess high levels of AlgT. Furthermore, we show that suppressors of toxic AlgT production have mutations in *mucP*, a protease involved in MucA degradation, and provide the first atomistic model of MucP. Our findings support a model where mutations in *mucP* stabilize the truncated form of MucA22 rendering it functional and therefore able to reduce toxicity by properly sequestering AlgT.

**IMPORTANCE:** *Pseudomonas aeruginosa* is an opportunistic bacterial pathogen capable of causing chronic lung infections. Phenotypes important for the long-term persistence and adaption to this unique lung ecosystem are largely regulated by the AlgT sigma factor. Chronic infection isolates often contain mutations in the anti-sigma factor *mucA* resulting in uncontrolled AlgT and continuous production of alginate, in addition to the expression of ~300 additional genes including *algT* itself. Here we report that in the absence of wild-type MucA, AlgT overproduction is lethal and that suppressors of toxic AlgT production have mutations in the MucA protease, MucP. Since AlgT contributes to the establishment of chronic infections, understanding how AlgT is regulated will provide vital information on how *P. aeruginosa* is capable of causing long-term infections.

## INTRODUCTION

Alginate production by the opportunistic pathogen *Pseudomonas aeruginosa* results in a mucoid phenotype and correlates with the establishment of a chronic infection (1–6). Production of alginate is highly regulated and is usually a response to external and internal membrane stressors (7–9). In nonmucoid strains, the anti-sigma factor MucA is inserted into the inner membrane where the cytosolic N-terminus interacts with the AlgT sigma factor (also called σ^22^ or AlgU) (10). AlgT is the master regulator of alginate biosynthesis genes, as well as about 300 other genes controlling phenotypes including pigment production, motility, and very long O antigen production (11–16). When AlgT is sequestered by MucA, the alginate biosynthesis operon is not transcribed and these strains produce little to no alginate.

AlgT is made available and alginate biosynthesis is triggered by a regulated intermembrane proteolysis (RIP) cascade that is activated by proteases (17). Briefly, activated AlgW cleaves the periplasmic C-terminus of MucA making the truncated inner membrane portion of MucA available for cleavage by MucP (18, 19). This releases the cytosolic N-terminus AlgT-bound fragment of MucA where it is finally completely degraded by the ClpPX proteasome. Once free of MucA, AlgT initiates transcription of its regulon (8, 20–22). AlgT itself is the first gene in an operon followed by *mucA, mucB, mucC,* and *mucD.* This operon is highly similar to the *rpoE, rseA, rseB,* and *rseC* operon in *Escherichia coli* (17, 23). AlgT and RpoE (also called σ^E^) are 66% similar and functionally interchangeable, with RpoE capable of complementing *algT* null mutants of *P. aeruginosa* (24). The upstream region of *algT* contains five promoters with two of these promoters themselves being AlgT-dependent (25, 26). Therefore, *algT* is positively autoregulated.

While nonmucoid strains are capable of producing and regulating alginate in certain environments, *P. aeruginosa* isolates from cystic fibrosis lung infections often overproduce alginate, termed the mucoid phenotype (1, 2, 27–29). The most common mutations that result in the mucoid phenotype are found in *mucA* (1, 30). The *mucA22* mutation, which is a deletion of a G in a string of 5 G’s, is the most common mutation observed in clinical isolates and results in a form of MucA with a truncated C-terminal domain (31). As a result, MucA22 is continuously subjected to RIP, AlgT is always free, and the alginate biosynthesis operon is always expressed (32, 33). Since AlgT promotes transcription of itself, mucoid strains have high levels of AlgT.

Despite the many discoveries surrounding the regulation of AlgT in nonmucoid strains, it is still unclear if mucoid strains are capable of regulating AlgT in the absence of wild-type MucA. Therefore, we sought to primarily study AlgT in the context of mucoid strain PDO300, a derivative of the nonmucoid laboratory strain PAO1 that contains the *mucA22* allele (31). However, we report here that the expression of exogenous *algT* is lethal in PDO300. Further, we show that suppressors of toxic AlgT production have mutations in *mucP,* a protease involved in MucA degradation. Our findings support a model where mutations in *mucP* stabilize the MucA22-AlgT complex ultimately rendering MucA22 functional. Since it has been reported that deletion of *mucA* is lethal (34), we propose that deletion of *mucA* is likely lethal due to deregulation and overproduction of AlgT.

## RESULTS AND DISCUSSION

### Overexpression of *algT* is lethal in *mucA22* strains

To study the expression of AlgT in mucoid *P. aeruginosa,* we began by making an IPTG-inducible copy of *algT* (P_tac_-*algT*) and inserted this construct in single copy at an ectopic site in PDO300. However, we found it difficult to grow PDO300 in the presence of inducer. Serial dilutions of PDO300 onto higher concentrations of IPTG revealed that increasing ectopic expression of *algT* reduced growth of PDO300 compared to growth without inducer (Fig. 1). Further, high levels of *algT* expression was entirely lethal (Fig. 1). This lethality was independent of alginate because growth of the nonmucoid strain PDO300 Δ*algD*, which has a clean deletion of the first gene in the alginate biosynthesis operon and does not produce alginate, was also inhibited by the expression of ectopic *algT*(Fig. 1). We also monitored growth of PDO300 and PDO300 Δ*algD* over time in liquid broth culture; without inducer, both strains grew to high density (Fig. S1). Conversely, when grown in the presence of inducer, PDO300 and PDO300 Δ*algD* had a decreased growth rate compared to uninduced conditions (Fig. S1), corroborating what we observed by serial plate dilutions (Fig. 1). Ultimately, both PDO300 and PDO300 Δ*algD* reached the same final density, likely due to the selection of mutants that can suppress toxic AlgT production. Altogether, these data suggest that in *mucA22* strains, such as PDO300, tightly control expression of *algT* is essential for growth.

**Figure 1.**
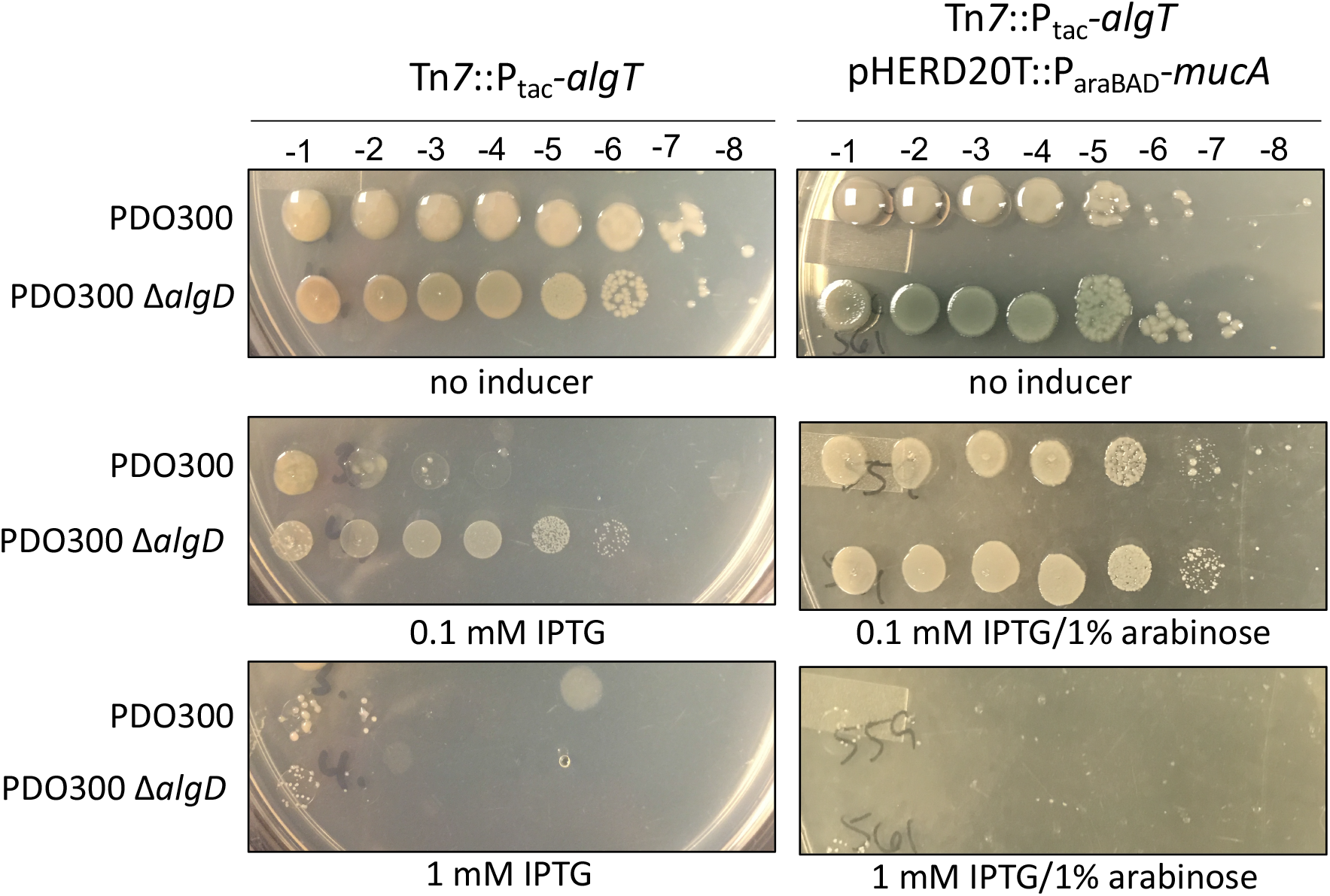
Overproduction of *algT* is lethal in *mucA22* strains and is partially rescued when wild type *mucA* is provided *in trans.* The *algT* coding sequence was cloned downstream of an IPTG inducible *tac* promoter and inserted, in single copy, at the Tn *7* site of each strain (Tn*7*::P_tac_-*algT*). Overnight cultures were grown without inducer, normalized to the same optical density, and then serially diluted onto LA containing no inducer, 0.1 mM IPTG, or 1 mM IPTG and/or 1% arabinose. IPTG induces expression of *algT* while arabinose induces expression of *mucA.* A) Strains were serially diluted onto increasing concentrations of IPTG to induce *algT* expression. Overexpression of *algT* inhibited growth and is independent of alginate production. Each strain on the right contains an arabinose-inducible copy of *mucA* on multicopy plasmid pHERD2OT (pHERD2OT::P_araBAD_-*mucA*). Corresponding dilution factors are shown on top. The strains shown are PAC539, PAC543, PAC559, and PAC561.

### Complementation with wild-type MucA partially rescues *algT* lethality

We hypothesized that *algT* lethality is due to the fact that PDO300 contains a *mucA* mutation. To investigate this, we monitored growth of PAO1 and PAO1 Δ*algD* (35, 36). PAO1 is the nonmucoid parent strain of PDO300 and contains wild-type *mucA.* When grown on lysogeny agar (LA), PAO1 is nonmucoid since *algT* expression is low. When the IPTG-inducible copy of *algT* is inserted into both strains and these strains grown on increasing concentrations of IPTG to induce *algT* expression, PAO1 became mucoid, as expected (37), while PAO1 Δ*algD* remains nonmucoid (Fig. S2). Both PAO1 and PAO1 Δ*algD* grew equally well when *algT* expression was induced indicating that in the presence of wild-type MucA, *algT* overexpression and accumulation is not lethal (Fig. S2).

If wild-type MucA is required to circumvent AlgT lethality, we hypothesized that expression of wild-type *mucA, in trans,* would rescue PDO300 from the toxic effects of *algT* overexpression. To test this, we expressed *mucA* from an arabinose-inducible plasmid and transferred this plasmid to the PDO300 and PDO300 Δ*algD* strains that contained the IPTG inducible promoter driving *algT.* On low levels of IPTG (to slightly express *algT)* and on high levels of arabinose (to overexpress *mucA*), growth of PDO300 and PDO300 Δ*algD* was no longer inhibited by the ectopic expression of *algT* (Fig. 1). Overexpression of *mucA* also rendered PDO300 nonmucoid, indicating that AlgT is being sequestered by MucA. Still, on higher levels of IPTG (1 mM), providing wild-type *mucA in trans* failed to rescue growth of either PDO300 or PDO300 Δ*algD* (Fig. 1). This is likely because not enough MucA is produced, even at this high level of induction, to sufficiently sequester AlgT to rescue these strains.

### Identification of mutations that inactivate AlgT

Over the course of our experiments we observed some PDO300 and PDO300 Δ*algD* colonies that were able to grow when *algT* was overexpressed. We isolated 8 suppressors (4 from PDO300 and 4 from PDO300 Δ*algD)* for further analysis. These strains grew on high levels of inducer and therefore were able to bypass the lethality caused by overexpression of *algT.* The most likely way to suppress *algT* toxicity would be to acquire mutations in one of the two copies of *algT.* To determine if this was the case, we sequenced both the native and inducible copies of *algT* and found that 2 out of the 8 suppressors had mutations in the cloned copy of *algT.* One had acquired a 9 bp duplication of nucleotides 136-144 (GAA GCC CAG) while the other had a C to T transition at nucleotide 400 that would result in a stop codon.

It has been observed that mucoid strains frequently revert to nonmucoid by acquiring mutations in *algT* (19, 25, 38). To determine what other mutations we could identify in *algT* that result in nonmucoid reversion, we grew mucoid PDO300 under low aeration, a stressful condition that selects for reversion (19). We isolated 18 PDO300 nonmucoid revertants and then sequenced *algT* to determine what proportion had acquired mutations in this gene. We found that 10 of the 18 (56%) of the nonmucoid revertants had mutations in *algT.* These data are similar to previous studies that estimated 41-83% of nonmucoid revertants contain *algT* mutations (14, 38, 39). We found that the same 9 bp nucleotide insertion as described above had occurred in 6 of our nonmucoid revertants. This insertion represented a duplication of nucleotides 127-135 (corresponding to amino acids DAQ) or nucleotides 136-144 (corresponding to amino acids EAQ), the insertion representing amino acids 43-45 or 46-48 (Fig. S3). Two studies have described a 9 bp insertion in this region, but whether this was a similar duplication was not indicated as the sequence of the insertion was not defined (27, 39).

How the duplication we observed affects AlgT is unclear since the insertion of 3 amino acids results in an in-frame mutation. To begin to assess how these mutations might be affect function, we constructed a model of AlgT in DNA-bound form (Fig. 2A, 2B, and S4) using a recent cryo-EM structure of the *E. coli* transcription initiation complex with RpoE (40). This model shows that domain I of AlgT comprising ~1-100 residues interacts with the −10 element of the promoter and plays crucial role in promoter melting. AlgT domain II binds to −35 element of the promoter. We also note that AlgT binds to MucA in a more compact structure (PBD 6IN7), wherein the domain I and II comes close (Fig. S4, AlgT conformation shown only). The flexibility of the 25-residue linker between the two domains might play a critical role in its promoter DNA binding. To determine how the 3 amino acid insertion alters AlgT domain I binding to DNA, we mapped the location of the DAQ and EAQ duplications to a highly conserved loop region in the AlgT model (Fig. 2C). To structurally interpret the effect of this duplication, we modeled the DAQ duplication in the loop region (Fig. 2D). The duplication distorts the promoter binding region and we also speculate that adding an additional negatively charged residue (Asp or Glu) further alters the binding strength between the loop and the crucial promoter melting region, further altering AlgT function and introducing a possible steric clash. Therefore, in this configuration AlgT would be unable to efficiently promote transcription, consistent with the nonmucoid reversion and loss of alginate production observed in these mutants. In *E. coli,* suppressors of high RpoE levels contain mutations in *rpoE,* but these are mostly near the −35 promoter-binding region and are proposed to weaken binding to promoters, but still support some transcription initiation (41), which is not what we observed for this *P. aeruginosa* mutant.

**Figure 2.**
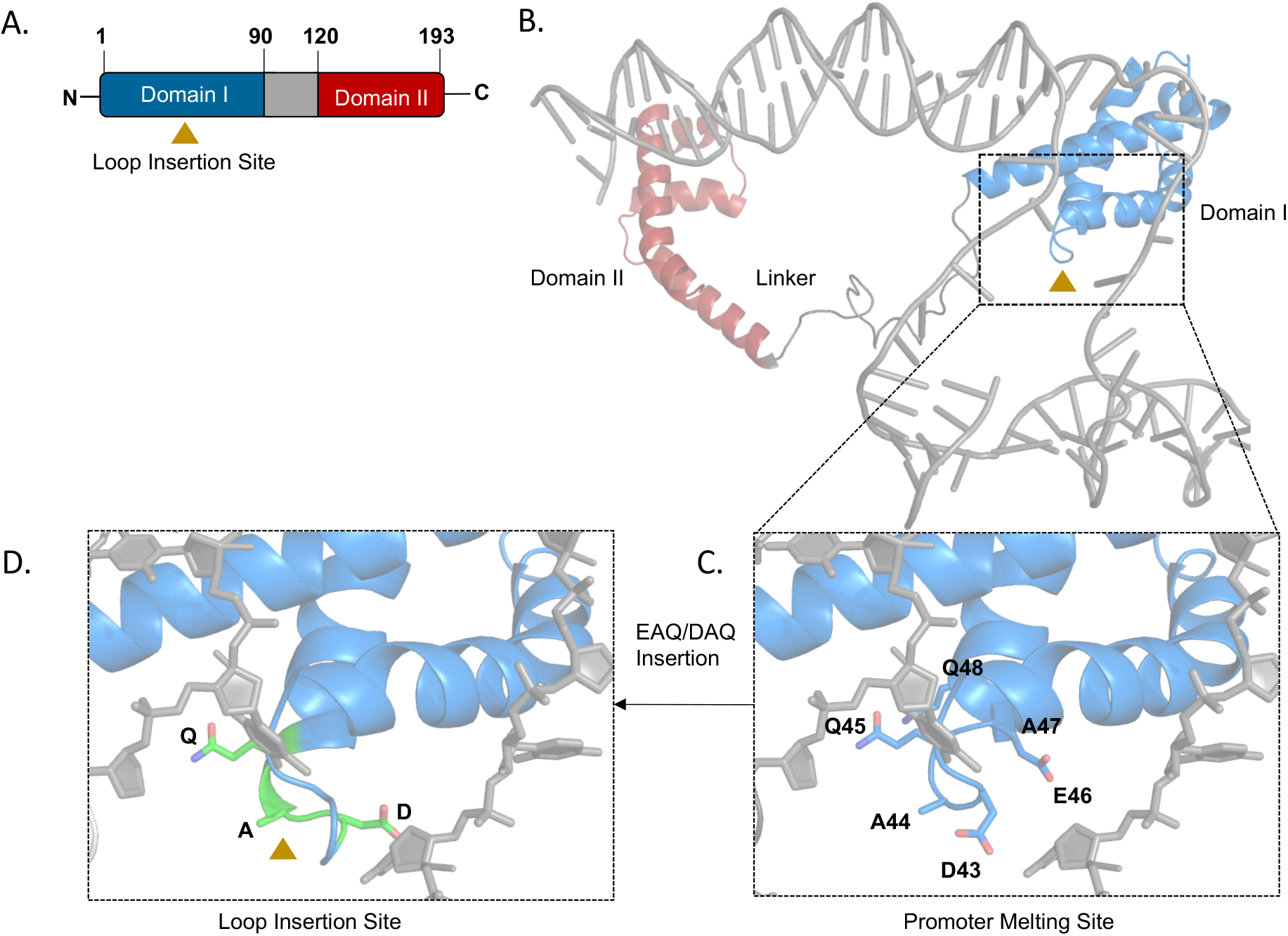
Model of AlgT bound to promoter DNA and the effect of the 3 amino acid insertion. A) Linear schematic of AlgT structure highlighting domain I (blue) and domain II (red), connected by a linker (grey). The loop insertion in the AlgT domain I occurs between D45 and E48 (golden triangle). B) Homology model of AlgT bound to promoter DNA modeled from RpoE sigma factor bound to DNA (from PDB 6JBQ). C) Residues in the DNA binding loop in domain I (D43 to Q48, shown as sticks) interact with DNA. D) The DAQ insertion (green) increases the size of the loop thus disrupting the optimal distance between the loop and the DNA, introducing steric clash.

### Suppressor mutations in *mucP* rescue AlgT-mediated toxicity

We performed whole genome sequencing to determine what mutations in the remaining 6 suppressor mutants were acquired that allowed for survival when *algT* was overexpressed. We found that 3 out of the 6 PDO300 and PDO300 Δ*algD* suppressors contained mutations in the MucA protease, *mucP.* MucP is a zinc metalloprotease that participates in the proteolytic cascade that degrades MucA (18, 19, 42). Delgado *et al.* analyzed the MucP sequence and found four possible transmembrane domains, one beta-loop domain, a metalloprotease zinc-binding motif, two PDZ binding domains, and a RIP motif (19). Many previous studies have predicted the secondary structure of the membrane bound region of MucP but failed to provide a three-dimensional atomistic model. The membrane bound region of MucP does not share overall sequence similarity to any protein for which a structure has been determined, so our initial trials for generating homology modeling or threading using the Swiss-Model, I-TASSER and LOMETS (43–45) servers were unsuccessful. Next, we attempted *ab initio* structure determination using QUARK (46, 47), which again failed to produce a model with conserved zinc binding domain. We next searched for distant homologs of MucP using InterPro database (IPR008915, www.ebi.ac.uk/interpro) and the resulting sequences were further sorted based on the zinc binding HXXXH and DGGH motifs using PROSITE database (https://prosite.expasy.org). Among these sequences, we found one homologous protease, from *Methanocaldococcus jannaschii,* for which the membrane bound structure was determined (PDB:3B4R). We used this model to generate a *de novo* MucP membrane bound structural model using helical constrains and active site geometry constrains from the HXXXH and DGGH motifs (Fig. S5A). As structural motifs are more conserved than sequence, M50 family proteases would be expected MucP to fold in a locally similar manner with the HXXXH and DGGH motifs, which coordinate a catalytically critical zinc ion, coming within 5Å distance. As structure determination of membrane proteins is difficult and the membrane region of MucP does not share any significant sequence similarity with known homologs for which structure is known, we used a combination of evolutionary and structural constrains and the partial template from *M. jannaschii* peptidase to generate the first atomistic model of *Pseudomonas* MucP.

The periplasmic region is composed of two PDZ domains which are connected by linkers to the membrane bound N-terminal and C-terminal domains of MucP (Fig. S5B). The membrane bound structure illustrates that any deletion, even in the C-terminal domain, would render the protein inactive as the complete catalytic site needs the ^21^HXXXH^25^ motif from N-terminal and the RIP motif ^403^DGGH^406^ from the C-terminal. Mutations destabilizing the helices of the MucP would also destabilize the fold and would inactive the protein.

One *mucP* suppressor mutant, found in PDO300, had an insertion of a “C” after nucleotide 358 (PDO300 *mucP*_358c_ins_) leading to truncation of the protein after 122 amino acids (Fig. 3). This truncates the protein before the first PDZ domain. The second *mucP* suppressor mutant was in PDO300 Δ*algD,* which had a deletion of nucleotides 910-916 (PDO300 Δ*algD mucP*_Δ910-916_), encoding “GCG GGG G”, leading to truncation of the protein after 316 amino acids (Fig. 3). Our model shows that this would truncate the protein just before the C-terminal membrane bound domain. Delgado *et al.* found the same 7 nucleotide deletion in MucP when looking for mutations that cause nonmucoid reversion (19). This mutation too, would make the peptidase function of MucP inactive due to loss of the ^403^DGGH^406^ RIP motif. The third *mucP* suppressor, also found in PDO300 Δ*algD,* had an in-frame deletion of nucleotides 1022-1036, encoding “CGC TCG ACT CCA TAA” (PDO300 Δ*algD mucP*_Δ1022-1036_), leading to early truncation of the protein after 445 amino acids (Fig. 3). This would again disrupt the catalytic site of MucP, making it inactive for its peptidase function. Overall, we predict each of these mutations interfere with the ability of MucP to degrade MucA22.

**Figure 3.**
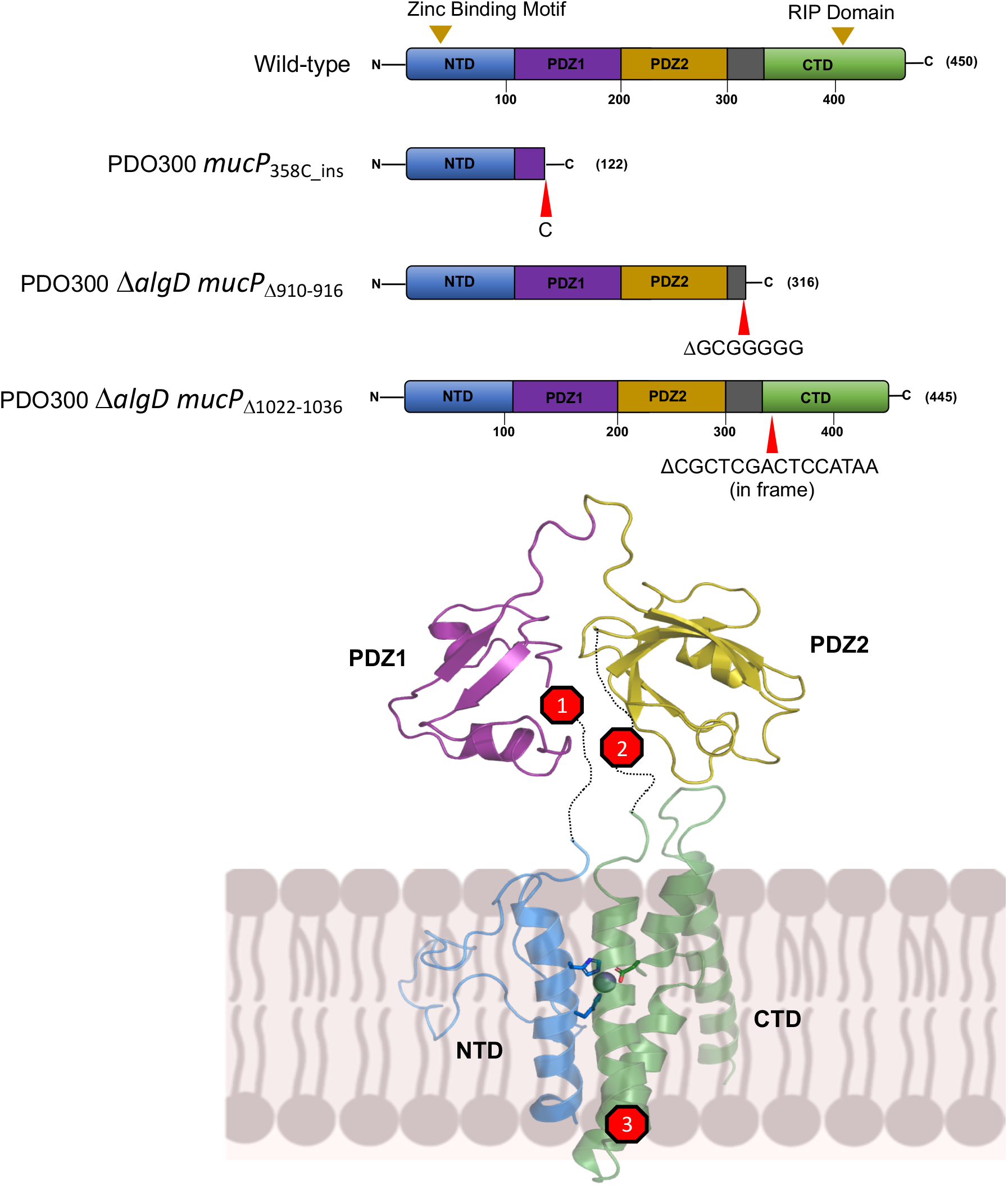
Suppressor mutations in *mucP* rescue AlgT-mediated toxicity. Depiction of the primary structure of wild-type MucP and suppressors (top). PDO300_538c_ins_ has an insertion of a “C” after nucleotide 355 leading to truncation of the protein after 122 amino acids. PDO300 Δ*algD mucP*_Δ910-916_ suppressor has a deletion of nucleotides 906-912, encoding “GGG GGC G”, leading to truncation of the protein after 316 amino acids. PDO300 Δ*algD mucP*_Δ1022-1036_ has an in-frame deletion of nucleotides 1019-1033, encoding “TAA CGC TCG ACT CCA”, leading to truncation of the protein after 445 amino acids. The location of each mutation is shown using a homology model of MucP (bottom). MucP, depicted in the inner membrane (IM), contains 4 transmembrane helices and a periplasmic region composed of two PDZ domains that are connected by linkers to the N-terminal (NTD) and C-terminal (CTD) membrane bound domain of MucP. The strains shown are PAC577, PAC579, and PAC582.

### Inactivation of MucP reduces expression of *algT*

We hypothesized that mutations in *mucP* would stabilize the MucA22-AlgT complex ultimately reducing toxic AlgT activity. If this is the case, we would expect that expression of genes controlled by AlgT would be reduced in *mucP* mutants. Since *algT* is autoregulated, we used an *algT* reporter as a readout of AlgT activity. To do this, the *algT* promoter region was transcriptionally fused to a *promoterless-lacZ* and then inserted, in single copy, at an ectopic site in each of the strains. In mucoid PDO300, *algT* promoter activity is ~2.5x higher compared to nonmucoid PAO1, which contains wild-type MucA (Fig. 4). When *algT* promoter activity is measured in the *mucP* suppressor mutant PDO300 *mucP*_358c_ins_, *algT* promoter activity is significantly decreased and comparable to PAO1 levels (Fig. 4). Overall, our results indicate that inactivation of MucP renders MucA22 functional and MucA22 is then able to sequester AlgT and reduce AlgT activity. As a result, *mucP* mutants bypass toxicity caused by the overexpression of *algT.*

**Figure 4.**
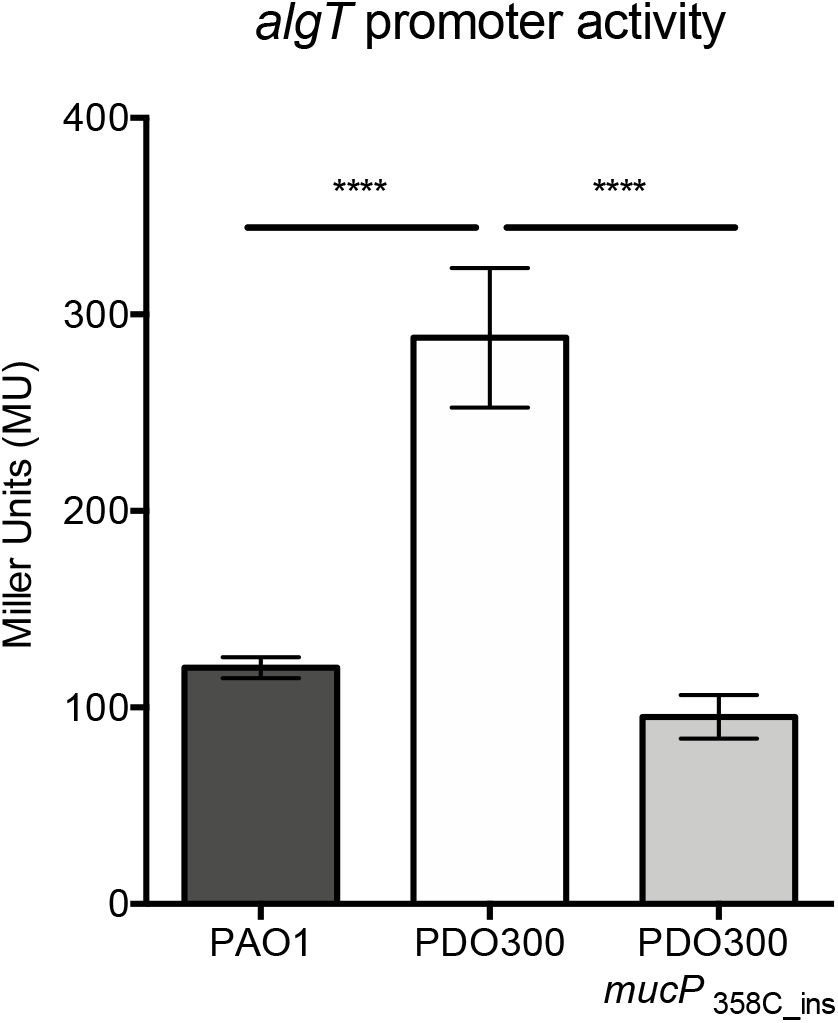
Inactivation of MucP reduces expression of *algT*. The *algT* promoter region was transcriptionally fused to a promoterless*-lacZ* to construct a *P5_aigT_-lacZ* reporter. The reporter was inserted at the CTX site of the *P. aeruginosa* chromosome of each of the strains. ß-galactosidase activity was determined during exponential phase growth in LB. Significance was determined using a one-way ANOVA with multiple comparisons. Error bars represent standard deviation (SD) of the mean from at least four biological replicates. **** P < 0.0001. The strains shown are PAC541, PAC539, and PAC577.

### Complementation of *mucP* suppressor mutants restores sensitivity to *algT* overexpression

Multiple attempts at constructing clean deletions of *mucP* in PAO1 and PDO300 were unsuccessful and therefore we could not test the hypothesis that deletion of *mucP* from PDO300 or PDO300 Δ*algD* would allow for growth when *algT* is overexpressed. In *E. coli,* the *mucP* homolog *rseP* is essential because the absence of RseP leads to loss of RpoE activity and the accumulation of unfolded outer membrane proteins (48, 49). In *P. aeruginosa,* however, *algT* does not appear to be essential under standard conditions as *algT* deletion strains have been reported (23, 50). If *mucP* is essential in *P. aeruginosa* and *algT* is not, this implies that MucP may have other targets, other than MucA-AlgT, that are critical for survival.

Since we could not delete *mucP,* we chose to overexpress *mucP* in the suppressor mutants PDO300 *mucP*_358c_ins_ and PDO300 Δ*algD mucP*Δ9io-9i6. We hypothesized that providing wild-type MucP *in trans* would render the suppressors sensitive to the overproduction of AlgT. To test this, we cloned the *mucP* gene on a multicopy arabinose-inducible plasmid and inserted this into both suppressor mutants. First, we confirmed that overexpression of *algT* in the presence of this plasmid was not lethal to the PDO300 and PDO300 Δ*algD* suppressors (Fig. 5). Likewise, overexpression of *mucP* by itself also did not inhibit growth of either strain. When *algT* and *mucP* are both induced, however, growth of both suppressor mutants was inhibited (Fig. 5). In summary, the suppressors were unable to grow due to complementation with wildtype *mucP* confirming that mutations in *mucP* are a mechanism to circumvent toxicity due to AlgT overexpression.

**Figure 5.**
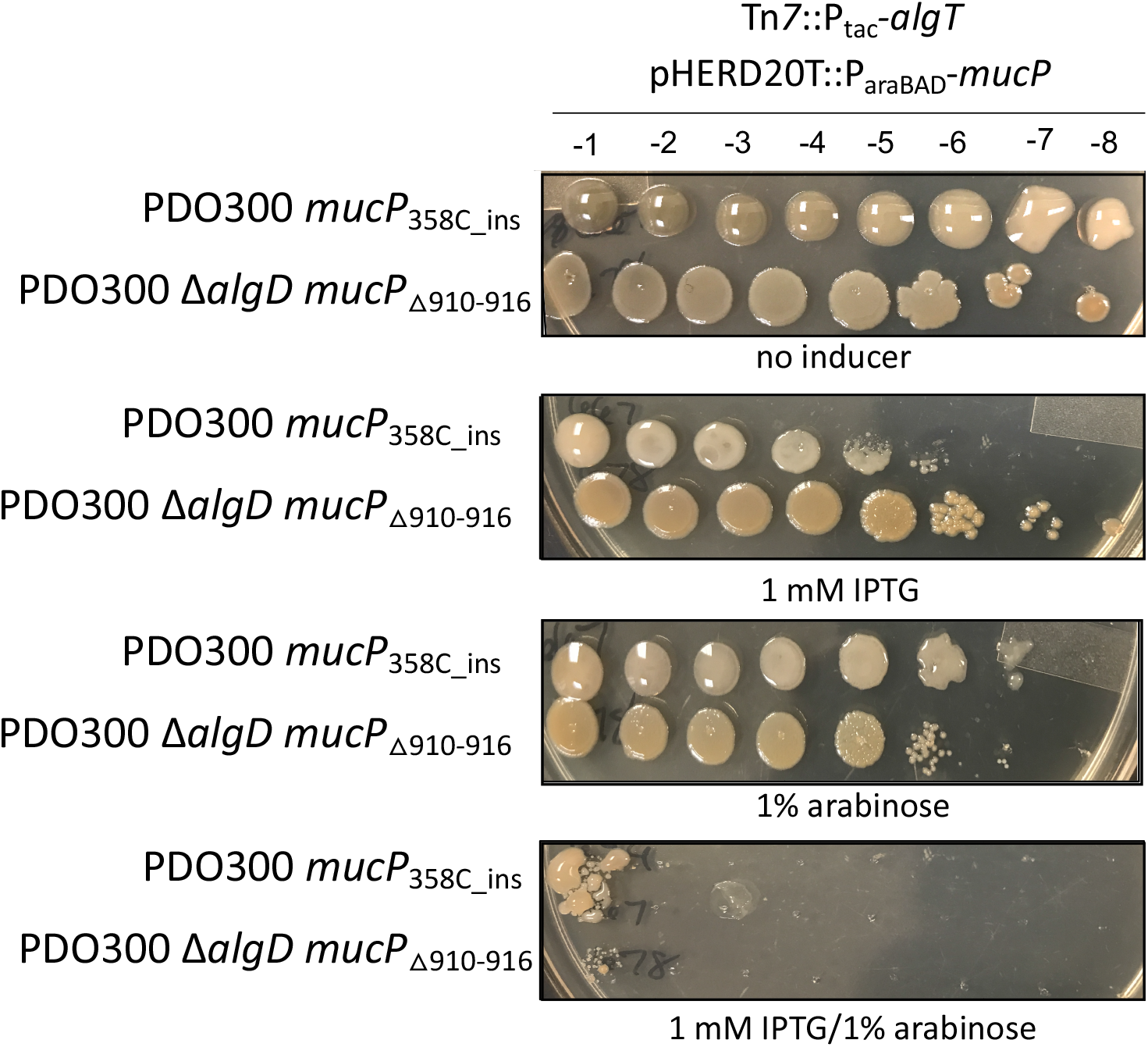
Complementation of *mucP* suppressor mutants restores sensitivity to *algT* overexpression. Strains were grown and as described in Figure 1. Each strain contains an IPTG inducible *algT* at the Tn7 site (Tn*7*::P_tac_-*algT*) and an inducible copy of *mucP* on multicopy plasmid pHERD2OT (pHERD2OT::P_araBAD_-*mucP*). IPTG induces expression of *algT* while arabinose induces expression of *mucP.* Corresponding dilutions factors are shown on top. Growth was only inhibited when both *algT* and *mucP* are overexpressed. The strains shown are PAC667 and PAC678.

### Conclusions

In mucoid strains containing a truncated form of the anti-sigma factor MucA, called MucA22, the cleavage site for the MucP protease is available and MucA22 is continuously degraded. As a result, MucA22 is no longer able to sequester the AlgT sigma factor and AlgT is available to direct transcription of its regulon, including the alginate biosynthesis operon. Here we show that overexpression and accumulation of AlgT in the presence of MucA22, but not wild-type MucA, is lethal. Suppressors of toxic AlgT contain mutations in the *mucA* protease *mucP*. When MucP is inactivated, we propose that MucA22 is not degraded and the truncated form of MucA22 is functional and able to sequester AlgT, reducing expression of the AlgT regulon and the subsequent downstream toxic effects.

It is well documented in *E. coli* that fine-tuning of RpoE is necessary to maintain cell envelope integrity (41, 51, 52). Deletion of *rseA* results in high RpoE activity leading to cell lysis in stationary phase (41, 53). Similarly, it is also not possible to delete *mucA* in *P. aeruginosa.* We propose that this is due to the toxic accumulation of AlgT. If the mechanism is similar to that in *E. coli,* this is hypothesized to be due to destabilization of the outer membrane. The *mucA22* mutation appears to bypass this toxicity over a *mucA* null mutation, likely because this form can be functional. Mutations in *mucP* are probably only one of the possible mechanisms for circumventing toxic AlgT accumulation. Future experiments aim to isolate and identify other suppressor mutants and to determine through what mechanism overproduction of AlgT leads to cell death.

## MATERIALS AND METHODS

### Bacterial strains and culture conditions

Bacteria were maintained on lysogeny agar (LA) containing 1.5% agar and grown in lysogeny broth (LB). When appropriate, media was supplemented with 15 μg/ml gentamycin, 10 μg/ml tetracycline, or 100 μg/ml carbenicillin for *Escherichia coli* and 60 μg/ml gentamycin, 100 μg/ml tetracycline, or 100 μg/ml carbenicillin for *P. aeruginosa.* Vogel-Bonner minimal medium (VBMM) plates supplemented with gentamicin and no salt LB (10 g/l tryptone and 5 g/l yeast extract) containing 15% sucrose were used for allelic exchange (54). Conjugations and sucrose counterselections were performed at 30°C. A list of strains, plasmids, and primers used are available in Table S1.

### Construction of PDO300 Δ*algD*

SM10 containing pEXG2-*mucA22* (14) was conjugated with PAO1 Δ*algD* following the puddle-mating protocol described by Hmelo *et al.* (54). After sucrose counterselection, single colonies were patched onto LA and LA containing gentamicin. Next, *mucA* was amplified, by single colony PCR, from gentamicin sensitive colonies using oAC089/oAC090 and sent to Genewiz for sequencing to identify colonies that contained the *mucA22* mutation.

### Construction of pHERD20T-*mncP*, miniCTX-*P5_algT_-optRBS-lacZ*, and generation of mutants

The *mucP* gene was amplified using oAC276/oAC277. This fragment was PCR purified (Qiagen) and ligated into HindIII digested gel-purified pHERD20T (37) using isothermal assembly (Gibson Assembly Master Mix, New England BioLabs). The reaction was transformed into chemically competent DH5α and selected for on carbenicillin. Plasmids were miniprepped (Qiagen) from carbenicillin resistant colonies and screened for the *mucP* insertion using oAC039/oAC040. To generate miniCTX-P5_*algT*_-optRBS-*lacZ*, 480 bps of the upstream *algT* region starting 20 bps upstream of the start codon, to incorporate the five *algT* promoters but not the native RBS, was amplified using oAC158/oAC159 and inserted into BamHI digested miniCTX1-optRBS-*lacZ* (14) using isothermal assembly (Gibson Assembly Master Mix, New England BioLabs). The reactions were transformed into chemically competent DH5α and selected for on tetracycline. Plasmids from single colonies were isolated (Qiagen Miniprep Kit) and screened for the *algT* promoter insertion using oAC32/oAC33. All plasmids in this study were electroporated into *P. aeruginosa* strains as previously described (55). miniTn*7*-P_tac_-*algT* (14) was selected for on gentamicin, miniCTX-P5_*algT*_-optRBS-*lacZ* was selected for on tetracycline, and pHERD20T-HA-*mucA* (23) and pHERD20T-*mucP* were selected for on carbenicillin.

### Isolation of nonmucoid revertants

A 1 ml culture of LB was inoculated with a single colony and incubated for 2-5 days, shaking vertically, at 37°C. Each day the culture was serially-diluted onto LB and incubated overnight at 37°C. Nonmucoid revertants were picked when the plate still contained both nonmucoid and mucoid colonies and therefore nonmucoid colonies could be compared to the mucoid ones. Using single colony PCR, *algT* were amplified using oAC107/oAC108, column purified (Qiagen miniprep kit), and sent to Genewiz for sanger sequencing.

### Isolation of suppressors, whole genome sequencing, and analysis

Overnight cultures of each suppressor strain grown without inducer were normalized to an OD_600_ of 0.5 and serially-diluted onto increasing concentrations of IPTG. Colonies that grew on high concentrations were considered suppressors of toxic *algT* expression. Genomic DNA was isolated using a DNeasy blood and tissue kit (Qiagen) and sent to the Microbial Genome Sequencing (MiGS) Center (migscenter.com). Genomes were quality trimmed using TrimGalore! v0.6.2 and reads having quality > 20 were assembled using SPAdes v3.13.1 (56, 57). Variant calling was performed using Snippy v4.4.0 (https://github.com/tseemann/snippy) using PAO1 as the reference genome (Assembly accession ASM676v1).

### β-galactosidase assays

Overnight cultures of each strain were back-diluted 1:100 into fresh LB and allowed to grow, rolling at 37°C, until exponential phase when samples were collected. Reporter assays were performed as previously described (58) with modifications (14).

### Computational methods for modeling of AlgT

Homology modeling of *Pseudomonas* AlgT was performed using Swiss Model (43) to generate a DNA bound conformation of the protein. The DNA bound structure of RpoE (PDB: 6JBQ) from *E. coli* was taken as the template to model a DNA bound conformation of AlgT. The model was further examined in Coot (59) to confirm favorable side-chain conformations near the DNA binding interface and minimize clashes, before a round of energy minimization in UCSF Chimera (60). The loop insertions ^43^DAQ^45^ and ^46^EAQ^48^ in AlgT were modeled in UCSF Chimera. We determined the sequence conservation of AlgT with the protocol described in Kuiper *et al.* 2019 (61). AlgT homologous sequences were retrieved by BLAST in UniProt. A set of ~200 unique amino acid sequences was then selected by applying a 90% sequence identity cut-off in CD-HIT (62).

### Computational methods for modeling of MucP

Calculations of site-specific conservation were made using Geneious Prime (63) for all MucP homolog sequences. *Pseudomonas* MucP has two regions i) membrane bound and ii) periplasmic, where the membrane bound M50 peptidase domain was predicted previously to be formed by both N-terminal (~1-100 residues) and C-terminal (~340-450 residues) sequences. NCBI BLAST searches identified two periplasmic PDZ domains, which were then modeled using Swiss Model with template PDB 2ZPL. The model was further examined in Coot (59) to confirm favorable side-chain conformations, before five rounds of energy minimization in UCSF Chimera (60).

## Supporting information

ACross_biorxiv2020_supplemental

## ACKNOWLEDGEMENTS

The authors would like to thank Graeme Conn for critical reading of the manuscript. ARC was supported by a predoctoral fellowship from the Cystic Fibrosis Foundation (CFF)-funded CF@LANTA RDP Center (MCCART15R0) and a NRSA predoctoral fellowship from the National Institutes of Health (NIH) under award number F31 AI136310. Additional funding was provided under award numbers NIH-R21AI122192 (JBG), CFF-GOLDBE16G0 (JBG), and CFF-DEY18F0 (DD). The content of the manuscript does not represent the official views of the NIH or CFF who had no role in study design, data collection, or interpretation of the data.

## AUTHOR CONTRIBUTIONS

ARC designed the experiments and wrote the paper; ARC, VR, ZW, and DD performed research and analyzed data; VR, ZW, DD, and JBG provided feedback on the experiments and the manuscript.

## DATA AVAILABILITY STATEMENT

The data that support the findings of this study are available from the corresponding author upon request.

