## Supplementary material for "Overproduction of the AlgT sigma factor is lethal to mucoid *Pseudomonas aeruginosa*": ACross_biorxiv2020_supplemental

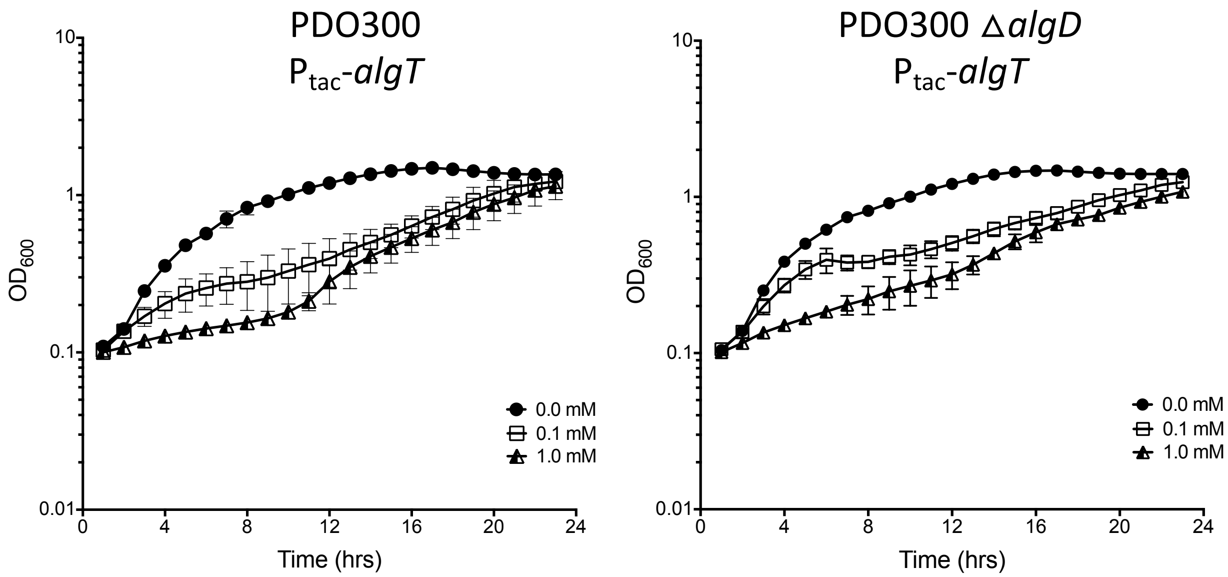


**Figure S1. Overexpression of *algT* in PDO300 and PDO300 Δ*algD* reduces growth.** Growth curves of each strain grown in LB or LB containing no inducer, 0.1 mM IPTG, or 1.0 mM IPTG. The strains shown are PAC539 and PAC543.

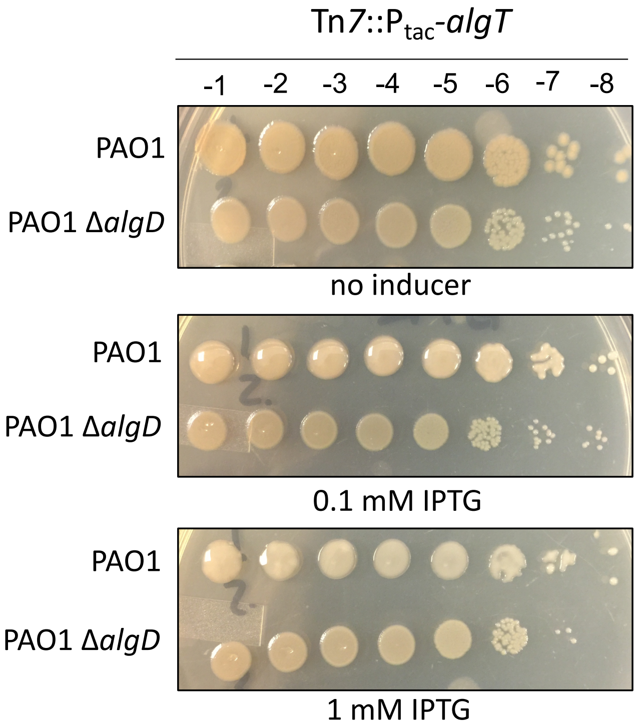


**Figure S2. Overexpression of *algT* in strains containing wild type MucA is not lethal.** The *algT* coding sequence was cloned downstream of an IPTG inducible tac promoter and inserted, in single copy, at the Tn*7* site of each strain (Tn*7*::P$_{tac}$-*algT*). Overnight cultures were grown without inducer, normalized to an optical density of 0.5, and then serially diluted onto LA containing no inducer, 0.1 mM IPTG, and 1 mM IPTG. IPTG induces expression of *algT.* Corresponding dilutions factors are shown on top. PAO1 becomes mucoid when *algT* is expressed. The strains shown are PAC501 and PAC541.

| | | | | | | | | | | | | | | | |
|---|---|---|---|---|---|---|---|---|---|---|---|---|---|---|---|
| | F | V | H | D | A | Q | | | | E | A | Q | D | V | A |
| ***algT*** | TTC | GTG | CAC | GAC | GCC | CAG | --- | --- | --- | GAA | GCC | CAG | GAC | GTA | GCG |
| | 117 | | | | | | | | | | | | | | 151 |
| | | | | | | | D | A | Q | | | | | | |
| ***dup1*** | TTC | GTG | CAC | GAC | GCC | CAG | GAC | GCC | CAG | GAA | GCC | CAG | GAC | GTA | GCG |
| | | | | | | | E | A | Q | | | | | | |
| ***dup2*** | TTC | GTG | CAC | GAC | GCC | CAG | GAA | GCC | CAG | GAA | GCC | CAG | GAC | GTA | GCG |

**Figure S3. AlgT sequence alignment of two nonmucoid revertants.** The AlgT amino acid sequence (green) is shown above the 117-151 bp nucleotide sequence (black). Both nonmucoid revertants had an in-frame duplication resulting in the insertion of three amino acids (red); DAQ (dup1) and EAQ (dup2). The underlined text represents the duplicated nucleotides.

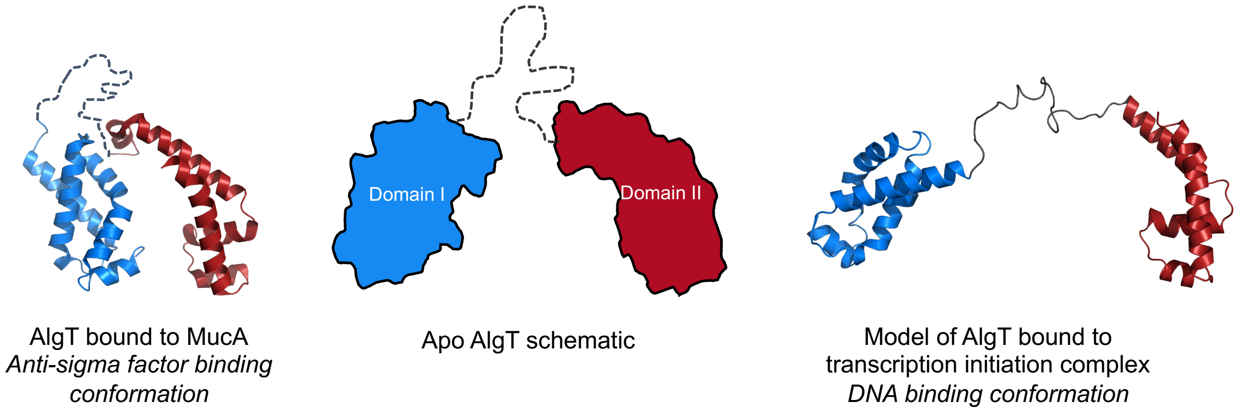


**Figure S4. Structural models of *Pseudomonas* AlgT.** The AlgT sigma factor is composed of two helical bundles which form the N-terminal domain I (blue) and C-terminal domain II (red) connected by a flexible 25 residue linker. The apo structure schematic is shown in the middle. Anti-sigma factor MucA binds AlgT and prevents it from binding to the RNA polymerase for transcription. The conformation of AlgT bound to MucA (PDB 6IN7) is shown in left, where domains I and II are arranged in a closed conformation. In contrast, AlgT interacts with the DNA in the transcription initiation complex with an extended linker conformation (right; modeled from RpoE-DNA complex, PDB 6JBQ), where domain I (in blue) engages the -10 element of the promoter and takes part in promoter melting, while the domain II (in red) interacts with -35 element of the promoter.

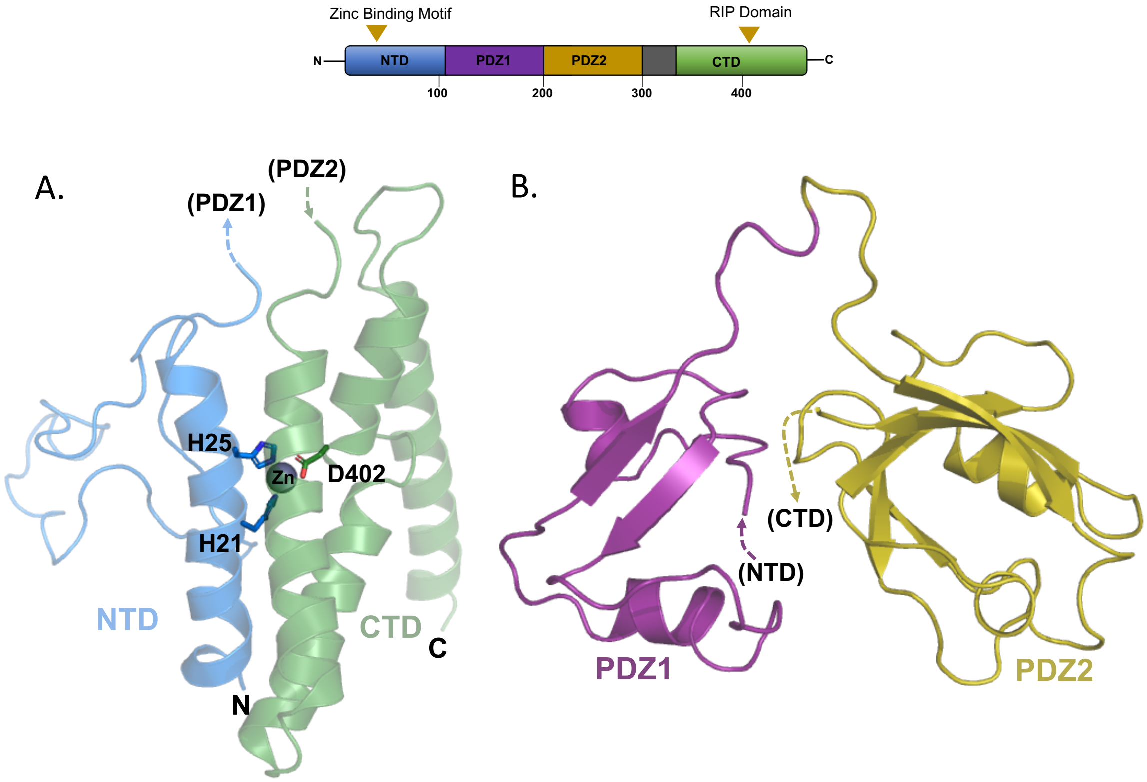


**Figure S5. Homology model of MucP membrane bound and periplasmic domains.** A) Model of membrane bound M50 peptidase domain of MucP modeled using *de novo* modeling using evolutionary and structural constrains with PDB 3B4R. As MucP does not share overall sequence similarity to any protein for which a structure has been determined, this model was generated *de novo* using structural constrains of the active site, helical constrains, secondary structure prediction, and using a distant homolog of MucP from *M. jannaschii*. B) Homology model of the two PDZ domains in MucP modeled with PDB 2FNE. PDZ1 is connected to the membrane bound N-terminal domain (NTD) while PDZ2 is connected to the membrane bound C-terminal domain (CTD). The catalytic core is composed of the NTD (His 21 and His 25), the CTD (D402), and a zinc metal cation.

**Table S1. Strains, plasmids, and primers used in this study.**

| Strains | Genotype or relevant features | Source |
|---|---|---|
| *E. coli* | | |
| DH5α | cloning background, plasmid maintenance | Invitrogen |
| *P. aeruginosa* | | |
| PAO1 | wild type (nonmucoid) | 35 |
| PDO300 | PAO1 *mucA22* (mucoid) | 31 |
| PAC342 | PAO1 Δ*algD* | 36 |
| PAC437 | PDO300 Δ*algD* | This study |
| PAC541 | PAO1 CTX::P5$_{algT}$-optRBS-*lacZ* Tn*7*::P$_{tac}$-*algT* | This study |
| PAC501 | PAC342 CTX::P5$_{algT}$-optRBS-*lacZ* Tn*7*::P$_{tac}$-*algT* | This study |
| PAC539 | PDO300 CTX::P5$_{algT}$-optRBS-*lacZ* Tn*7*::P$_{tac}$-*algT* | This study |
| PAC543 | PAC437 CTX::P5$_{algT}$-optRBS-*lacZ* Tn*7*::P$_{tac}$-*algT* | This study |
| PAC559 | PAC539 pHERD20T-HA-*mucA* | This study |
| PAC561 | PAC543 pHERD20T-HA-*mucA* | This study |
| PAC667 | PAC577 pHERD20T-*mucP* | This study |
| PAC678 | PAC579 pHERD20T-*mucP* | This study |
| PAC577 | PAC539 suppressor (*mucP* 358 C insertion) | This study |
| PAC578 | PAC539 suppressor (*algT* duplication 136-144 GAAGCCCAG) | This study |
| PAC579 | PAC543 suppressor (*mucP* deletion 910-916 GCGGGGG) | This study |
| PAC581 | PAC543 suppressor (*algT* C400T) | This study |
| PAC582 | PAC543 suppressor (*mucP* deletion 1022-1036 CGCTCGACTCCATAA) | This study |
| **Plasmids** | **Description** | **Source** |
| miniTn*7*-P$_{tac}$-*algT* | IPTG inducible *algT* in single copy | 14 |
| pHERD20T | arabinose-inducible multicopy plasmid | 37 |
| pHERD20T::P$_{araBAD}$-*mucA* | arabinose inducible *mucA* with an N-terminus HA tag | 23 |
| pHERD20T::P$_{araBAD}$-*mucP* | arabinose inducible *mucP*, multicopy | This study |
| pEXG2-*mucA22* | *mucA22* allelic replacement vector | 14 |
| miniCTX-optRBS-*lacZ* | promoterless-*lacZ* reporter with optimized RBS, single-copy | 14 |
| miniCTX-P5$_{algT}$-optRBS-*lacZ* | *algT* promoter-*lacZ* reporter | This study |

| **Primers** | **Sequence** | **Source** |
|---|---|---|
| oAC039 | ATCGCAACTCTCTACTGTTTCT | 37 |
| oAC040 | TGCAAGGCGATTAAGTTGGGT | 37 |
| oAC089 | TTCCACACATTATACGAGCCGGAAGCATAAAT GTAAAGCAatgagtcgtgaagccctgca | 14 |
| oAC090 | CGAGCTCGAGCCCGGGGATCCTCTAGAGTCGA CCTGCAGAtcagcggttttccaggctgg | 14 |
| oAC107 | TGAGCCCGATGCAATCCAT | This study |
| oAC108 | CAACTGGTAACGCGACCAG | This study |
| oAC158 | CTGGAGCTCCACCGCGGTGGCGGCCGCTCTAG AACTAGTGatgcgcaggtgttccggaag | This study |
| oAC159 | AATCATGGTCATAGCTGTTCCTCCTTACTGCAG CCCGGGGggaggagcttcgagcgtccc | This study |
| oAC276 | GGTACCCGGGGATCCTCTAGAGTCGACCTGCA GGCATGCAatgagtgcgctttacatgat | This study |
| oAC277 | TTTTCCCAGTCACGACGTTGTAAAACGACGGC CAGTGCCActacagacgactcagatcgt | This study |

*lowercase nucleotides anneal to plasmid sequence during isothermal assembly
